# Remote underwater photographs reveal environmental correlations and patterns in reef manta ray habitat use in Laamu Atoll, Maldives

**DOI:** 10.64898/2026.05.09.723939

**Authors:** Benjamin J. Guilford-Pearce, Miriam Staiger, Guy M. W. Stevens, Philip D. Doherty, Jinaad Ali

## Abstract

Reef manta rays (*Mobula alfredi*) are threatened by fishing and other anthropogenic threats. Which, when coupled with conservative life history traits, have made this species vulnerable to extinction. Spatiotemporal ecological knowledge, such as site fidelity and visitation patterns to key aggregation sites, are imperative for effective conservation management of *M. alfredi*. A novel method of environmental sensing, remote underwater photo systems (RUPs), was employed to understand drivers of *M. alfredi* habitat use and resighting patterns. RUPs were deployed at four cleaning sites around Laamu Atoll, Maldives. Between March 2021 and May 2023, 455,458 photos were analysed. Generalised linear models revealed increases in *M. alfredi* presence in response to high chlorophyll-a concentrations, low illumination moon states, the Southwest Monsoon, and in the morning, while human presence had no effect. Branchial spot patterns allowed for 81 *M. alfredi* individuals to be identified, from 629 sightings, representing 51.59% of Laamu Atoll’s previously identified population (n = 157). Cleaning stations are visited more intensively during periods of increased productivity of the Southwest Monsoon, likely in response to greater foraging opportunities near the study areas. Additionally, moon state, used as a proxy for tidal strength, was associated with increased visitation during new moon periods, suggesting that weaker tidal states may facilitate presence. These data support integrating RUPs with observational surveys to improve inferences about habitat use and our understanding of cleaning sites frequented by *M. alfredi*. This study aims to inform the implementation of Laamu Atoll’s first marine protected area management plan.

## Introduction

The world’s ocean ecosystems support a great abundance and diversity of life, but have faced increased pressure from anthropogenic exploitation since the Industrial Revolution [1]. Scientific research has increasingly highlighted declines in oceanic and neritic elasmobranch populations [2,3], none more so than ray species inhabiting the shallow waters of tropical ecosystems, such as coral reefs [2]. Thirty-two percent of elasmobranchs are now threatened with extinction, and a further 37.5% lack sufficient data to make assessment possible [2–4]. Indeed, the reef manta ray (*Mobula alfredi*), which is found in the Indo-West Pacific on shallow coral reefs throughout tropical and subtropical waters, is listed as Vulnerable to extinction on the IUCN’s Red List of Threatened Species [5]. Recent studies call for legal protective measures to be accompanied by regionally specific, community-engaged, and holistic management, as current marine protections are generally inadequate to fully protect *M. alfredi* and their habitats [6–9].

Globally, M. alfredi populations are fragmented and range in size from fewer than 100 to many thousands of individuals, with population connectedness non-linear. Geographical barriers, such as deepwater channels and remote archipelagos, as well as consistent and sufficient sources of food fuelling strong site fidelity, serve as key regulators of *M. alfredi* dispersal [10–12]. These conspicuous and charismatic zooplanktivores have evolved for a life of continuous motion and spend much of their time in waters shallower than 50 m, searching for productive foraging patches to sustain their large body mass [13–15]. However, emerging research demonstrates the role that *M. alfredi* plays in connecting offshore and inshore waters and in foraging on both pelagic and benthic zooplankton [16,17]. *Mobula alfredi* is considered an extreme *K*-selective species, with a gestation length of over a year, long postpartum recovery periods, late maturation ages, single-pup pregnancies, and slow growth rates [7,9,18]. These life-history characteristics, coupled with relatively small and fragmented populations and aggregatory behaviours, increase the impact of fishing pressure and have led to the over-exploitation and decline of manta ray populations globally. Collectively, targeted capture for meat and gill plates, unintentional bycatch in active and ghost-fishing nets, boat strikes, habitat degradation, climate change-related effects, and unmanaged tourism activities threaten *M. alfredi* [19–22]. For example, between 2010 and 2016, Baa Atoll, Maldives, welcomed more than 25,000 tourists annually, which has been shown to affect *M. alfredi* behaviours in 37% of encounters [20]; implied to cumulatively negatively impact fitness [23]. Recent declines in *M. alfredi* populations are threatening their ecological and socioeconomic importance, with less developed countries where *M. alfredi* populations persist displaying high levels of economic dependence on the species [15,24–27]. Marine protected areas (MPAs) are an essential tool in the spatiotemporal protection of specific areas of importance, such as cleaning sites and habitats of highly mobile marine vertebrates, like *M. alfredi*, but their success varies [6,28]. Considering this, few of these important aggregation sites in the Western Indian Ocean are fully protected from fishing, and less than 0.1% of the Maldives’ exclusive economic zone is recognised as Important Sharks and Ray Areas [29].

The intrinsic vulnerability of *M. alfredi* coupled with their threats, requires adequate MPAs to conserve their ecological and socio-economic importance. However, without effective management plans and active enforcement thereof, these MPAs often provide little to no meaningful conservation benefit and fall into the ‘paper park’ paradigm, a ubiquitous issue in ∼70% of MPAs globally [6,30–32]. There are anecdotal reports of successful management of *M. alfredi* [20,33]. For example, Hanifaru MPA in Baa Atoll, Maldives, exemplifies how *M. alfredi* aggregations, if managed consistently, can mitigate threats such as negative human-manta interactions [20]. However, until the last decade, limited understanding of the threats and species ecology, a lack of research effort, and a notable absence of commercial interest have led to *M. alfredi* being understudied and ultimately under-protected [29,34,35]. To aid protective efforts, there is a need for specific ecosystem-based knowledge (Grorud-Colvert et al. 2021; Stewart et al. 2018). Marine protected areas at *M. alfredi* aggregations can provide substantial benefits via relatively small, informed areas of protection [28,36]. While regions like the Chagos archipelago can feasibly protect entire local *M. alfredi* populations’ home ranges [8], often, this is not socioeconomically or politically feasible [36,37]. Therefore, a baseline of ecological knowledge of a site’s use must be developed to enable marine spatial planners to effectively allocate limited resources for *M. alfredi* conservation across their entire range.

In the Maldives, human underwater and fisheries observations have historically been used to develop region-specific baselines of knowledge [7,38]. Although these methodologies are extremely useful for a variety of research topics, such as population demographics, reproductive ecology and mark-recapture studies, they do have considerable limitations. These include the possible effect of human presence on behaviour [20] and the logistical restrictions which in-person surveying often imposes on effort [14]. Remote underwater photo (RUP) systems have proven effective for manta ray population assessments, and long-term, unbiased, and continuous monitoring of cleaning sites [10,39]. Visiting sites to be cleaned is an important daily function for many large reef vertebrates [40–42]. *Mobula alfredi* are no exception, regularly aggregating at these cleaning sites to get cleaned, engage in reproductive behaviours, avoid predators, and to thermoregulate [7,43–45]. Much smaller in size when compared to key foraging areas, cleaning sites are easier to monitor continuously [46–48]. Therefore, applying this RUP method to monitor *M. alfredi* visitations to cleaning sites is a useful tool for quantifying drivers of site use and ultimately prioritising protective management recommendations.

The Maldives has the world’s largest known population of *M. alfredi* [7], with tourism estimated to generate ∼US$311 million annually [49], a 380% increase since 2008 [50]. *Mobula alfredi* within the Maldives are nationally protected, and their direct capture is illegal [51], although their unintentional capture by fishers does occur [21]. Merely 0.5% of the Maldivian exclusive economic zone (EEZ) is protected by 42 MPAs, with the aim to increase this to 30% by 2030 [19]. Only one MPA has an active management plan, delineating most MPAs within the Maldives as little more than paper parks [19,52,53]. In the Maldives, bycatch, habitat degradation, boat strikes, and unmanaged tourism are key threats to this population [7,21,54]. Maldives-wide findings have highlighted the nation’s overall importance, but many have struggled to focus on the more remote southern atolls of Huvadhoo, Fuvahmulah, Addu, and Laamu due to logistical constraints [7,38,54]. Effective protection depends on a detailed understanding of local populations and their habitats [43,55]. For example, knowledge of *M. alfredi* has informed the establishment of MPAs in the Maldives, including Hanifaru and Anga Faru [56]. Without atoll-specific studies, conservation measures risk being ad hoc and misinformed.

This study aims to further our understanding of Laamu Atoll’s *M. alfredi* population by assessing their spatiotemporal use of cleaning sites via passive RUP systems placed at four key aggregation sites within the atoll. Specifically, we aim to assess which environmental variables drive presence at these sites, what re-sightings of individuals reveal about their residency patterns and movement behaviour, and demonstrate the effectiveness of RUPs.

## Materials and methods

### Study site

The Maldives, situated in the central Indian Ocean, comprises 26 geographical atolls (Fig 1). Laamu Atoll consists of 82 small coral islands (69 of which are uninhabited), broken by only six channels. The atoll is large, covering 48 km north to south and 35 km east to west (Fig 1). Encircled by abyssal water [57], Laamu Atoll experiences biannual, monsoon-driven upwelling of nutrient-rich waters [54]. The Maldives Manta Conservation Programme has consistently studied Laamu Atoll’s *M. alfredi* population since 2014 and has identified key areas of habitat use by this species. RUPs were deployed at four known *M. alfredi* cleaning sites: Hithadhoo Corner, Fushi Kandu, Boduhuraa Beyru and Fonadhoo Beyru (Fig 1, Table 1).

**Fig 1.**
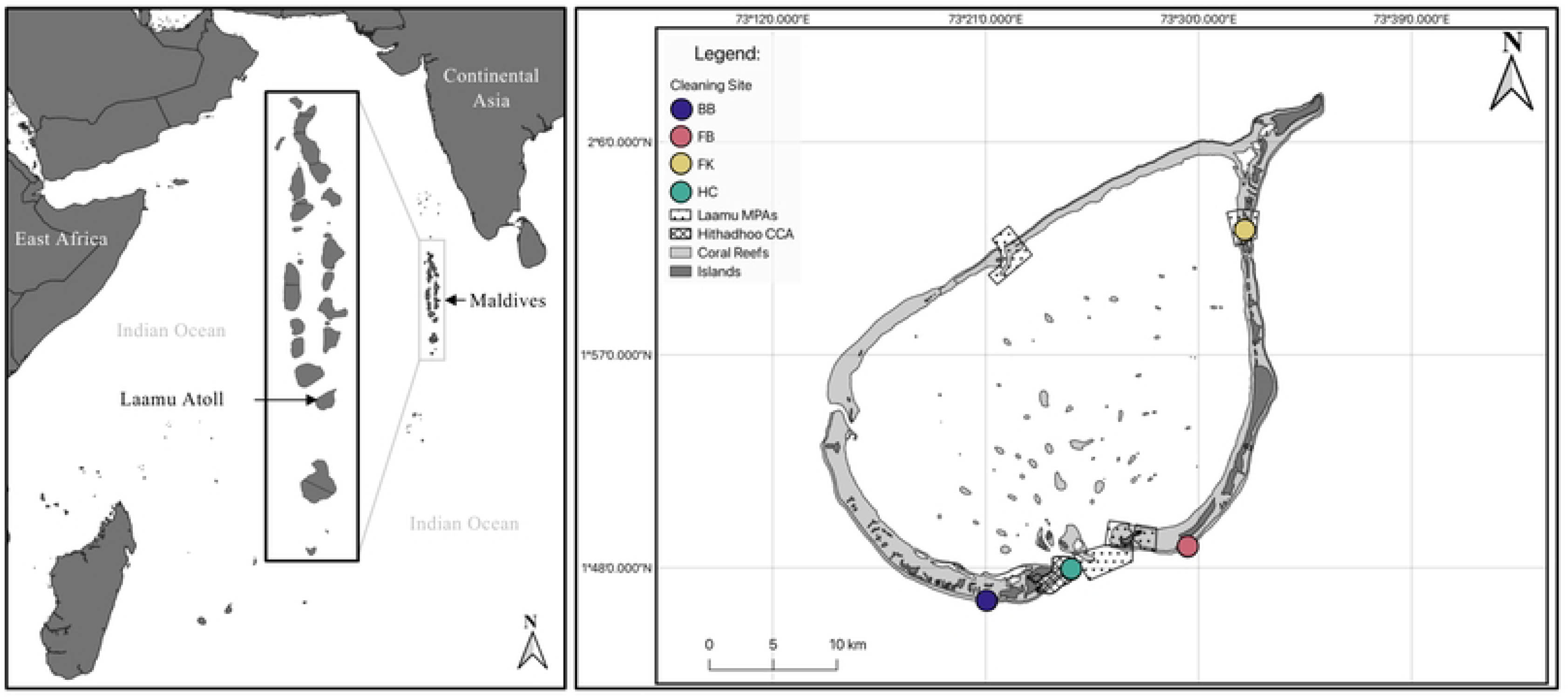
Study location. **A)** Location of the Republic of Maldives in the Indian Ocean and enlargement of the 26 geographical atolls. **B)** Map of Laamu Atoll with the four remote underwater photo (RUP) deployment locations: Boduhuraa Beyru (BB; Purple), Fonadhoo Beyru (FB; Red), Fushi Kandu (FK; Yellow) and Hithadhoo Corner (HC; Green). Polygons detailing designated marine protected areas (MPA) and Hithadhoo Baaneykolhu Community Conservation Area (CCA), recently proposed by the Hithadhoo Council.

**Table 1.** Summary of survey sites.

| Name | Co-ordinates | Observed period | Location type | Depth (m) | Size (m) |
| --- | --- | --- | --- | --- | --- |
| Hithadhoo Corner | 1°47'53.76"N<br>73°24'35.99"E | 29/04/2021<br>-<br>18/05/2023 | Channel | Shallow Block: 14.5 – 17.7m<br>Yellow: 18.3 – 22.2m | Shallow Block: 24x12m*<br>Yellow: 17x18m |
| Fonadhoo Beyru | 1°48'52.21"N<br>73°29'32.03"E | 13/09/2022<br>-<br>20/03/2023 | Outer-reef Slope | 24.9 – 26.7m | 5.5x6.5m |
| Fushi Kandu | 2° 2'19.12"N<br>73°31'57.29"E | 08/05/2022<br>-<br>21/04/2023 | Channel | 11.4 – 15.3m | 13x8m* |
| Boduhuraa Beyru | 1°46'35.57"N<br>73°21'4.78"E | 05/01/2022<br>-<br>13/05/2023 | Outer-reef Slope | Deep: 26 – 28m*<br>New: 23 – 25m* | Deep: 10x10m*<br>New: 8x5m* |

\*Denotes an estimated value.

### RUP deployment

All sites were accessed from Six Senses Laamu Resort on Olhuveli Island (Fig 1) by boat using SCUBA. Hithadhoo Corner is at the western edge of the Gaadhoo-Hithadhoo Channel (Fig 1) and consists of five distinct coral bommie cleaning station blocks: Shallow Block, Ridge, Yellow, Split and Turtle, sampled from April 2021 until May 2023 (S1 Fig). RUPs were primarily deployed on Shallow Block due to frequent observations of *M. alfredi* cleaning during a trial phase. Before July 2021, RUPs were placed at Yellow, Split and Turtle for a collective total of 23 days. Fushi Kandu is a single cleaning site, with RUPs deployed from May 2022 until April 2023. Boduhuraa Beyru is an outer reef site characterised by several cleaning locations, two of which were used for RUP deployments from January 2022 to May 2023 (Table 1). Fonadhoo Beyru is another outer-reef site where RUPs were deployed at a single deep-cleaning station from September 2022 to March 2023. Research at all sites was granted by the Maldives Government (Protected Species Research Permit: 2021: EPA/2021/PSR-M07; 2022: EPA/2022/PSR-M07; 2023: EPA/2023/PSR-M07; Marine Scientific Research Permits in the Maldives: 2021: (OTHR)30-D/INDIV/2021/70; 2022: (OTHR)30-D/INDIV/2022/9; 2023: (FRM)30-D/PRIV/2022/54).

Three RUP systems were used: a single GoPro HERO® 4 (12 megapixels, wide-angle mode, 128GB memory) and two GoPro HERO® 8 Black cameras (12 megapixels, wide-angle mode, 128GB memory), all with ∼160° fields of view [58]. Cameras were connected to Voltaic Systems^©^ batteries (19,200 mAh), and time-lapse settings were configured to capture a single-frame photo every minute during daylight hours. Deployments were typically scheduled between 05:59 and 17:59, although soak times varied. Cameras were secured in custom-built, waterproof housings with standardised orientations. Shallow Block had a custom-made concrete block placed on the bare reef to attach RUPs. A compass was used at the Fushi Kandu, Boduhuraa Beyru, and Fonadhoo Beyru sites to ensure orientation continuity. Supplementary straps and weights were used to secure the cameras onto dead or bare reef, and to optimise the camera’s view. Rubble was used to camouflage the systems.

#### RUP photo analysis

The maximum number of *M. alfredi* and/or humans observed in the frame (maxN) was recorded for each photo. Upon a sighting of any part of either *M. alfredi* or humans in the frame, an estimation of the total period a specific site was used was summarised into sighting events. Sighting events were considered terminated if either had not been seen for 10 minutes. Sightings of a single photo were classed as one minute in length. Sighting durations were defined as the interval between the first photo of a sighting and the last. The maximum maxN for a sighting event was used as a proxy for the total number of *M. alfredi* for that specific event and binned into the nearest hour (±), ranging from 05:00 to 19:00. An estimate of the relative abundance of *M. alfredi* and humans using a specific cleaning site over the day was the summed maxN of that day’s sighting events.

Adequate photos of branchial spot patterns were identified, as per Marshall and Pierce [59], and recorded as a confirmed number of *M. alfredi*. They were manually and automatically matched to the regional Laamu Atoll database using trained researchers and the Manta Trust’s image-recognition matching tool IDtheManta (https://mantabase.org/home). Until project termination, 157 unique *M. alfredi* have been identified in Laamu Atoll, and >6,000 in the Maldives as a whole [60].

### Environmental variables

#### Daily

Environmental variables were measured to understand their influence on *M. alfredi* abundance. Large seasonal changes lack definite and consistent start and end times [61]. Despite this, seasonality was defined as described by Anderson *et al.* [38]; May to October, the Southwest (SW) Monsoon, December to March, the Northeast (NE) Monsoon, and April and November as the two months of monsoonal retreat. Moon phases were extracted from an online database (https://www.timeanddate.com) and categorised into four states: new, first quarter, full, and third quarter and assigned to each survey day. Tidal charts were obtained from the Maldives Meteorological Service (https://www.meteorology.gov.mv) for Gan, Laamu Atoll and used for all cleaning sites. Tide states were defined into four categories: high, ebb, low, and flood. High and low tides were rounded to the nearest hour and categorised into three hours: the hour of and ±1 hour [42]. Interval periods between low and high, and between high and low, were defined as the flood and ebb, respectively. Both consisted of three hours but varied with tidal periods. Daily 0.25° x 0.25° chlorophyll-a (Chl-a; mg/m^-3^) data, from depths of ∼0.4 m, were obtained from Copernicus (https://www.copernicus.eu) through the Global Ocean Biogeochemistry Analysis and Forecast product [62] and assigned to each survey day. These data were sourced from models whose results are highly correlated with satellite and BGC-Argo measurements (correlation coefficient: 0.81, RMSD: 0.59, [63]). Water temperature was measured every minute at Shallow Block, Hithadhoo Corner, from July 2021 to April 2023 using HOBO^®^ Pro-V2 loggers (±0.2 °C) and TCM-x Current Meters (Lowell Instruments LLC, ±0.1 °C) and the data were averaged into daily bins. HOBO^®^ loggers were placed adjacent to the RUP system deployment location and TCM-x meter was strategically placed to reduce physical current obstructions from the cleaning station (S1 Fig).

#### Diurnal

The identification of fine-scale oceanographic factors which affect *M. alfredi’s* presence at shallow reef habitats [64] elicited the use of current meters over an hourly period rather than daily. The TCM-x Current Meter uses 3-axis accelerometers and magnetometers to obtain current speed (cm/s) and heading (0 - 360 °). It was set to record every minute between December 2022 and March 2023 at Shallow Block. Speed and heading were averaged into hourly bins.

### Statistical Analysis

Sighting event maxN, summarised into the daily estimated number of *M. alfredi* and sum hourly maxN, provided the foundations of relative abundance in this study. Relationships between estimated abundance and observed counts (maxN and confirmed) were assessed using quasi-Poisson generalised linear models (GLM) with a log link to account for overdispersion. Spearman’s rank correlations were additionally used to assess monotonic associations between variables. Significant positive correlations support the effectiveness and inclusion of the estimated abundance of *M. alfredi* as a response variable.

Generalised linear models and Generalised Linear Mixed Models (GLMMs) were used to assess the effects of environmental variables on *M. alfredi* abundance at four cleaning stations around Laamu Atoll, using the ‘*stats*’ and ‘*lme4*’ [65] packages in R, respectively [66]. Two response variables derived from the same sighting event maxN data were analysed here to understand seasonal and diel drivers of abundance, respectively: one, the estimated *M. alfredi* abundance counted in daily intervals (GLM 1.1 and 1.2) and two, the maxN summarised into hourly ones (GLMM 1.1 and 1.2).

GLM 1.1 consisted of a negative binomial distribution with the daily estimated *M. alfredi* abundance as the response and the environmental variables: Chl-a, moon state, cleaning station and monsoon season as fixed predictors. A Poisson distribution could not be used due to overdispersion. An interactive term was included between monsoon and cleaning station to account for the nuanced influence of the monsoon across different sides of the atoll [54]. To compensate for varying deployment lengths per day and, therefore, the number of photos available for analysis, an offset for the log of the number of photos per day was also specified in the model. In total, 317 days and three cleaning sites (Fushi Kandu, Boduhuraa Beyru and Fonadhoo Beyru) lacked temperature recordings. Therefore, GLM 1.2 was a subset of the daily variables, including only the days and sites with temperature recordings, and was modelled using a negative binomial distribution with the following fixed predictors: Chl-a, moon state, monsoon season, and temperature. Again, a Poisson model could not be used due to overdispersion, and the same offset was modelled as GLM 1.1.

A Poisson-distributed GLMM with a log link function was used to understand the diurnal abundance of *M. alfredi*. GLMM 1.1 used the sum of sighting-event maxN in hourly bins as a response variable, with the fixed predictors: sum of hourly maxN humans, hour of day, cleaning station, and tide state. GLMM 1.2 looked to include current speed and heading as predictors. Due to only being recorded at Hithadhoo Corner and over three months, a subset of data was used to construct a Poisson-distributed GLMM with a log link function with the sum hourly maxN of *M. alfredi* as the response and the fixed predictors: sum hourly maxN of humans, time of day, tide state, current heading and current speed. A random effect for Julian Day was included in both GLMMs to account for daily variation in cleaning station use.

The dredge function from the ‘*MuMIn*’ package in R [67] was used to compare all possible combinations of explanatory variables for GLM 1.1 and 1.2 and GLMM 1.1 and 1.2, separately. Throughout model simplification, the offset and random effect were fixed for GLM 1.1 and 1.2 and GLMM 1.1 and 1.2, respectively. Model simplification used two asymptotically bias-corrected parsimony [68]: corrected Akaike Information Criterion (AIC_c_) and Bayesian Information Criterion (BIC). Absolute values of AIC_c_ and BIC were not used for assessment, rather a comparison of the differences over other candidate models (ΔAICc and ΔBIC, Burnham & Anderson 2002), calculated by the following, where *i* is the model:

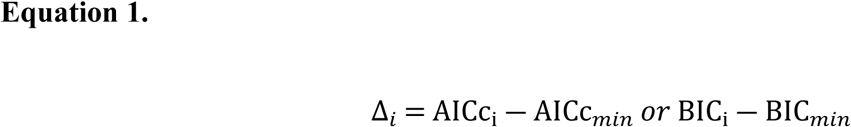

The relative quality of a model was assessed using the criteria presented by Burnham and Anderson (2002), whereby the model with the lowest value (either AIC_c_ or BIC) has the greatest support and therefore a Δ*_i_* of 0. Models with Δ*_i_* values <2 were used as the threshold, with models with Δ_i_ values>2 AIC considered inadequate for interpretation and disregarded from further analysis [69]. To assess the relative model fit of GLMs with Δ*_i_* values of <2, McFadden’s pseudo-*R^2^* (*ρ^2^*) was calculated via the R package ‘*pscl*’ [70]. McFadden’s pseudo-*R^2^* compares the log-likelihood of a candidate model of the null. Outputs range from 0, indicating no improvement over the null, to 1, indicating a perfectly fitted (undesirable) saturated model [71]. As high values of *ρ^2^* (>0.4) suggest over-fitting, McFadden [71] notes that those between 0.2 and 0.4 are indicative of an excellent fit. Additionally, two pseudo-*R^2^* estimates (tri-gamma) were used to provide an understanding of GLMM model fit using the ‘*MuMIn*’ package in R [67]. These include the conditional-*R^2^* (*R^2^_GLMM(c)_*), which is interpreted as the variance explained by both fixed and random effects and marginal-*R^2^* (*R^2^ _GLMM(m)_*), which only considers the fixed effects [72].

## Results

### Overview

A total of 132 RUP systems were deployed between April 2021 and May 2023 (S2 Fig). These deployments spanned 736 days, resulting in a combined duration of 7,579 hours and 41 minutes, and a total of 455,458 photos analysed, of which 9,414 (2.06%) contained *M. alfredi*. The greatest proportion of images with *M. alfredi* present in a single day was 36.57% (n = 192) on 5^th^ October 2022 at Hithadhoo Corner throughout a deployment from 09:16 until 18:00 (524 minutes). The median sighting duration was 4 minutes (IQR: 1.25 – 7.27, range: 1 – 74). Additionally, the largest number of *M. alfredi* individuals present over one day was 22, at Hithadhoo Corner on the 16^th^ of January 2022. The clarity of photos varied due to biofouling, floating debris and reef fish obstructing the view (S3 Fig). Logistical constraints lead to inconsistency in survey effort between sites (S2 Fig). From all photos analysed, Hithadhoo Corner consisted of 70.36% (n = 320,466), compared to 16.76% (n = 76,354) at Fushi Kandu, 7.24% (n = 32,977) at Boduhuraa Beyru and 4.68% (n = 21,303) at Fonadhoo Beyru.

### Daily Patterns

Sighting events estimated a total of 2,600 *M. alfredi* sighted over the study period, with a mean daily sighting of 3.53 ± 4.38 *M. alfredi* (mean ± SD; range 0 – 22). Daily sightings varied, whereby 240 days had no estimated *M. alfredi,* and 87 had ≥ 10. *M. alfredi* sightings also varied spatially, with Hithadhoo Corner and Fushi Kandu having similar mean estimates per photo of 0.58 ± 0.68% (n = 1870; range = 0 - 3.05%) and 0.52 ± 0.66% (n = 403; range = 0 - 2.75%), respectively. Fonadhoo Beyru displayed a higher mean estimated occurrence of *M. alfredi* per photo, with 0.81 ± 0.64% (n = 173; range 0 - 2.4%) and Boduhuraa Beyru a lower mean estimated occurrence of 0.27 ± 0.39% (n = 91; range= 0 - 1.89%). Estimated abundance was a significant positive predictor of both maxN and confirmed counts in quasi-Poisson generalized linear models (maxN: β = 0.107 ± 0.004 SE, *p* = <0.001; confirmed: β = 0.150 ± 0.008 SE, *p* = <0.001), corresponding to multiplicative increases of 1.11 and 1.16 in expected counts per unit increase in estimated abundance, respectively. These patterns were consistent with Spearman’s rank correlations, indicating strong monotonic associations between estimated abundance and maxN (ρ = 0.86, *p* = < 0.01) and confirmed counts (ρ = 0.66, *p* = < 0.01, S4 Fig).

The full GLM 1.1 model explained 17.62% of the deviance in *M. alfredi* sightings and had a pseudo-R^2^ value of 0.071, indicating a moderate model fit (Table 2). Based on AIC_c_ and BIC, the full model was the top-ranked model, and no other model fell within two AIC_c_ or BIC units of it (Table 2). The difference between the full and null models was large (ΔAIC_c_ = 1061.36; ΔBIC = 993.09; *ρ^2^* = 0.046; Table 2), indicating that the modelled predictors increase our understanding of the variance in *M. alfredi* cleaning site abundance. The low deviance suggests unmodelled parameters might be affecting the identified variance in presence. The abundance of *M. alfredi* at cleaning sites was positively associated with Chl-a concentration (Fig 2). Furthermore, moon phase influenced *M. alfredi* abundance, with early phases within the lunar cycle, especially the first quarter, showing higher abundances at cleaning sites than later moon phases (Fig 3).

**Fig 2.**
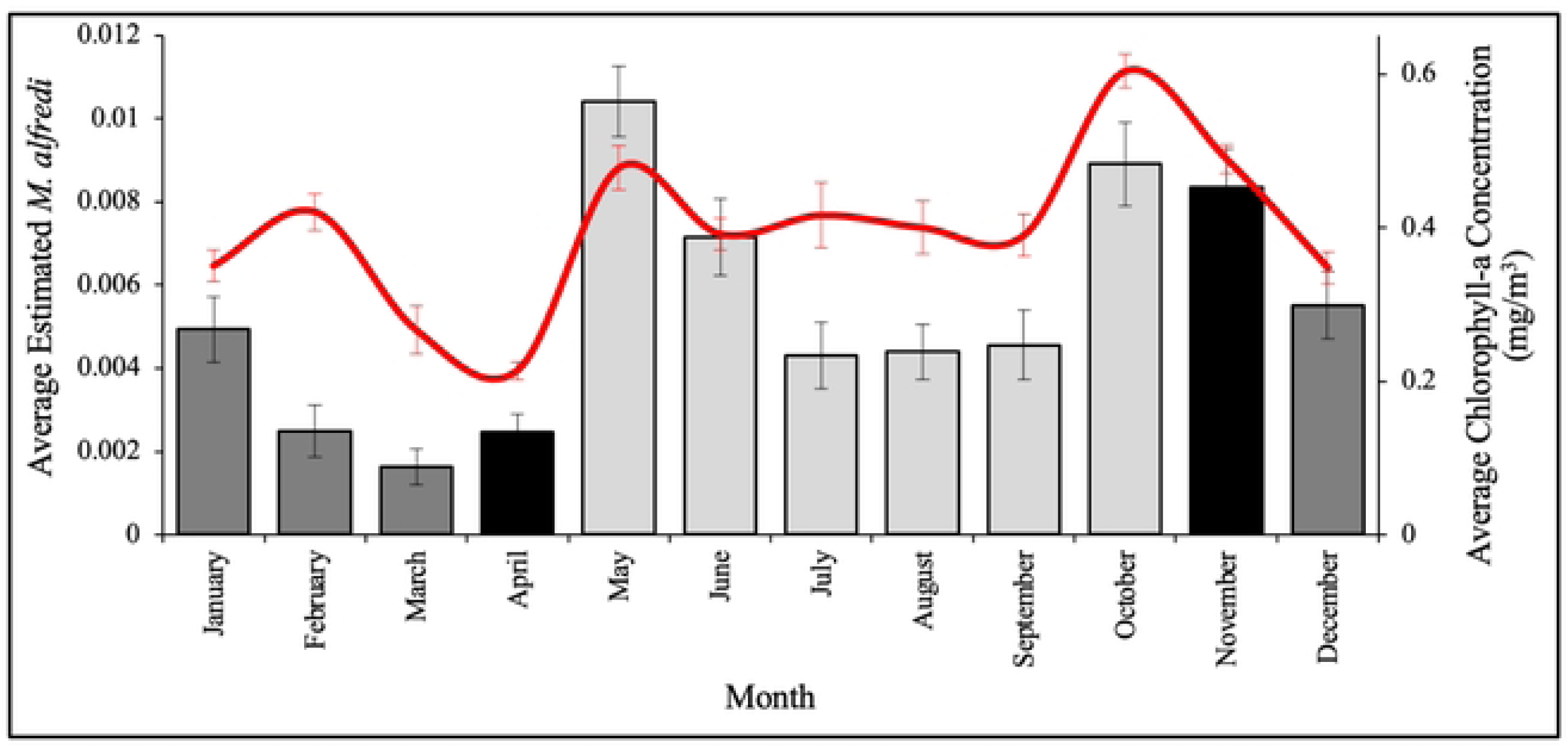
Seasonal *M. alfredi* presence. Average estimated (∑MaxN sighted / day) *Mobula alfredi* per photo (bar) and average chlorophyll-a concentration across Laamu Atoll, Maldives (mg/m3; red line) for each month across all studied sites around Laamu Atoll. Shading represents the monsoon season; light grey: Southwest Monsoon, Grey: Northeast Monsoon and Black: monsoonal transition.

**Fig 3.**
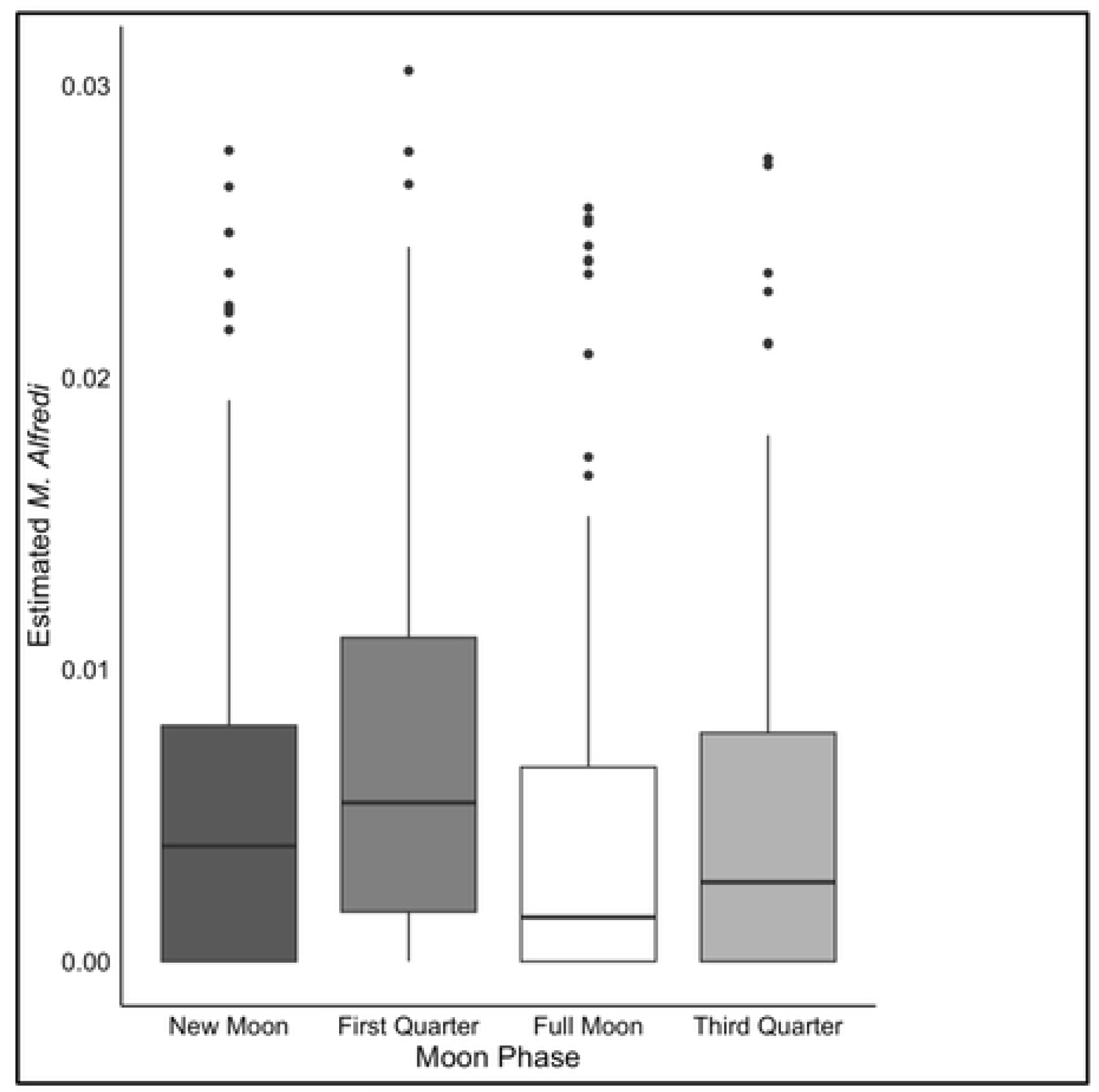
*Mobula alfredi* cleaning site presence during different moon phases. The effect of the moon state on the estimated (∑MaxN sighted / day) *Mobula alfredi* per photo. The box denotes the interquartile range, the line the median and the line to the minimum and maximum values within 1.5 times the interquartile range from the upper and lower quartiles, respectively.

**Table 2.** Model Summaries.

| Model | Response | Predictors | Offset | df | Loglik | AICc | ΔAICc | BIC | ΔBIC | Model Fit ($\rho^2$ and $R^2$) | DE(%) |
| --- | --- | --- | --- | --- | --- | --- | --- | --- | --- | --- | --- |
| GLM 1.1 - Negative Binomial | Daily manta maxN | ~ chla + moon + site * season | log(number of photos) | 16 | -10644.21 | 21321.2 | 0 | 21394 | 0 | $\rho^2 = 0.071$ | 17.62 |
| | | ~ 1 | | 1 | -11190.26 | 22382.5 | 1061.36 | 22387.1 | 993.09 | $\rho^2 = 0.025$ | 0 |
| GLM 1.2 - Negative Binomial | Daily manta maxN | ~ chla + moon + season | log(number of photos) | 7 | -6043.78 | 12101.8 | 0 | 12130 | 0 | $\rho^2 = 0.056$ | 12.93 |
| | | ~ chla + moon + season + temp | | 8 | -6043.65 | 12103.7 | 1.82 | 12135.8 | 5.8 | $\rho^2 = 0.056$ | 12.94 |
| | | ~ 1 | | 1 | -6261.58 | 12525.2 | 423.34 | 12529.2 | 399.25 | $\rho^2 = 0.023$ | 0 |
| GLMM 2.1 - Poisson | Hourly total manta maxN | ~ time + site + (1 \| Julian Day) | N/A | 19 | -5161.28 | 10360.7 | 0 | 10494.5 | 0 | $R^2_{\text{GLMM(C; M)}} = 0.47; 0.40$ | 7.05 |
| | | ~ time + site + human + (1 \| Julian Day) | | 20 | -5161.26 | 10362.6 | 1.97 | 10503.5 | 9.01 | $R^2_{\text{GLMM(C; M)}} = 0.47; 0.41$ | 7.05 |
| | | ~ 1 | | 2 | -5552.76 | 11109.5 | 748.85 | 11123.6 | 629.12 | $R^2_{\text{GLMM(C; M)}} = \text{N/A}$ | 0 |
| GLMM 2.2 - Poisson | Hourly total manta maxN | ~ time + (1 \| Julian Day) | N/A | 16 | -292.76 | 618.6 | 0 | 685.6 | 37.25 | $R^2_{\text{GLMM(C; M)}} = 0.67; 0.61$ | 7.79 |
| | | ~ time + heading + (1 \| Julian Day) | | 17 | -291.72 | 618.7 | 0.07 | 689.8 | 41.43 | $R^2_{\text{GLMM(C; M)}} = 0.69; 0.63$ | 8.11 |
| | | ~ time + human + (1 \| Julian Day) | | 17 | -292.53 | 620.3 | 1.68 | 691.4 | 43.04 | $R^2_{\text{GLMM(C; M)}} = 0.68; 0.62$ | 7.8 |
| | | ~ time + heading + human + (1 \| Julian Day) | | 18 | -291.51 | 620.4 | 1.8 | 695.6 | 47.27 | $R^2_{\text{GLMM(C; M)}} = 0.68; 0.63$ | 8.3 |
| | | ~ 1 + (1 \| Julian Day) | | 2 | -317.91 | 639.8 | 21.25 | 648.3 | 0 | $R^2_{\text{GLMM(C; M)}} = \text{N/A}$ | 0 |

Models within either two $\Delta\text{AICc}$ or $\Delta\text{BIC}$ units of the top-performing model and the null model ( $\sim 1$ ) for all four responses tested. Model
statistics shown include: the degrees of freedom (df), the logarithm of the likelihood function (Loglik), the AICc and BIC test statistics and
their differences ( $\Delta$ ), metrics of model fit which include a McFadden pseudo- $R^2$ ( $\rho^2$ ) for the GLMs and a conditional (C) and marginal (M)
tri-gamma pseudo- $R^2$ ( $R^2_{\text{GLMM(C; M)}}$ ) for the GLMMs and the percentage deviance explained (DE).

The retention of the interaction term between the cleaning site and monsoon indicates that cleaning site abundance varies with the monsoon (Fig 4). The Southwest Monsoon and the retreat had higher mean abundances of *M. alfredi* per photo (0.0069 ± 0.0071; 0.0051 ± 0.006, respectively) than the Northeast Monsoon (0.0037 ± 0.0056, Fig 4). Fushi Kandu appeared to be the cleaning site most affected by the monsoon, with a large increase in abundance during the Southwest Monsoon and a considerable drop during the retreat and Northeast Monsoon (Fig 4). Alternatively, Hithadhoo Corner was the site least affected by the monsoon, with *M. alfredi* cleaning year-round (Fig 4). Fonadhoo Beyru and Boduhuraa Beyru were surveyed for only 32 and 61 days, respectively. Fig 4 shows a reduction in *M. alfredi* cleaning-site use at Boduhuraa Beyru, especially during the Southwest Monsoon, potentially indicating *that M. alfredi* uses this area opportunistically.

**Fig 4.**
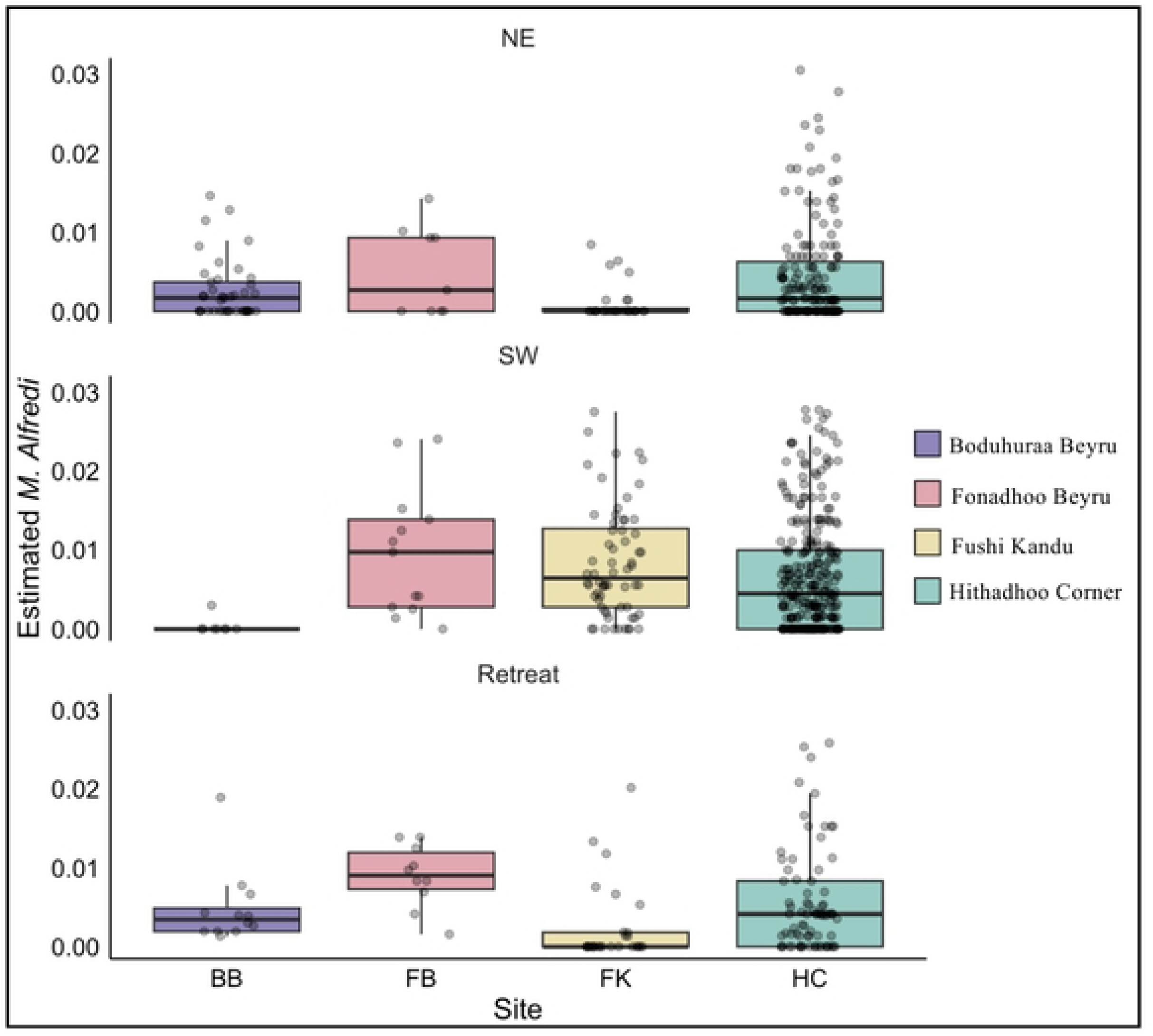
Cleaning site-specific differences in *Mobula alfredi* presence. The average estimated (∑MaxN sighted / day) Mobula alfredi per photo, and the interaction between the monsoonal season and the cleaning stations. The box denotes the interquartile range, the line the median and the line to the minimum and maximum values within 1.5 times the interquartile range from the upper and lower quartiles, respectively.

The inclusion of temperature in GLM 1.2 resulted in a negligible 0.01% increase in the deviance explained by the model, despite its status as a predictor excluded from the top model (Table 2). Furthermore, temperature contributed to modest increases in both the AICc and BIC, with the AICc indicating strong support and the BIC substantially less support. This discrepancy suggests that the increased complexity introduced by temperature does not sufficiently improve model fit and has no discernible effect on *M. alfredi* abundance after accounting for other predictors.

### Diurnal Patterns

Diurnal variation in *M. alfredi* abundance was analysed to understand temporal drivers of daytime abundance. Overall, 80.1% of hours had a sum maxN of 0 and 16.5% had a maxN of ≥ 2. These varied spatially, with Hithadhoo Corner having its highest maxN of 4, and a mean maxN of 0.3 ± 0.69 per hour (n = 6,047). Fushi Kandu, Fonadhoo Beyru, and Boduhuraa Beyru had peak maxN and mean maxN per hour of 4 and 0.29 ± 0.76; 3 and 0.45 ± 0.81; and 5 and 0.14 ± 0.46, respectively (n = 1416, 1809, and 647). Boduhuraa Beyru expressed the highest recorded maxN of 5 *M. alfredi* in a single photo, and Fushi Kandu recorded the highest sum maxN per hour of 8.

The top-performing model for the Poisson-distributed GLMM 2.1 included time and cleaning station, omitting tide state and human abundance (Table 2). This model explained 7.05% of the deviance and had an R^2^_GLMN (C, M)_ of 0.47, 0.4. Between 8:00 and 13:00, *M. alfredi* cleaning site abundance was highest (Fig 5). Although there was a slight increase in abundance during the flood (0.35 ± 0.77) and low tide (0.31 ± 0.74), the difference between the ebb (0.24 ± 0.59) and high tide (0.29 ± 0.69) was marginal.

**Fig 5.**
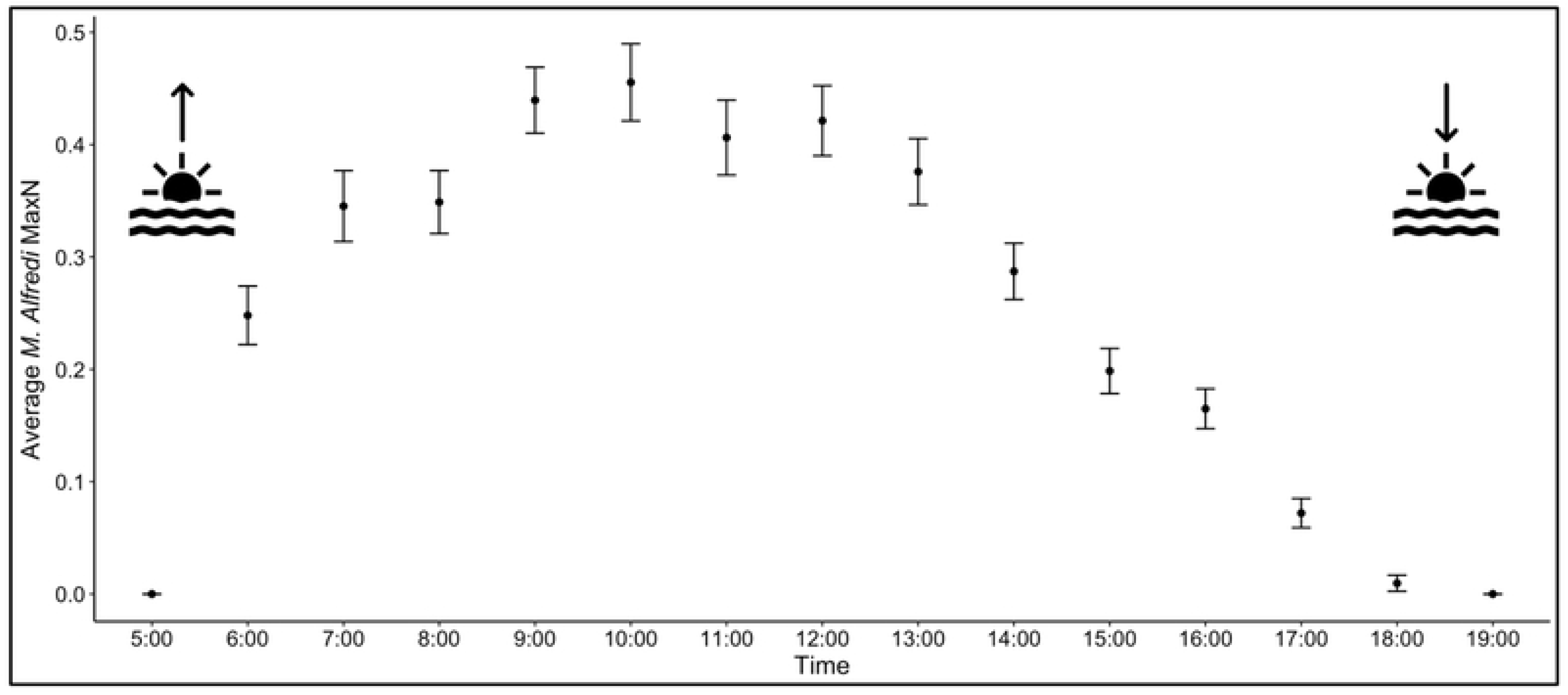
Diurnal cleaning site presence. The mean and standard error (±) maxN of sighted *Mobula alfredi* for each hour interval at all sites.

One other model from GLMM 2.1, which included human abundance, was within two AIC_c_ units of the top-performing model and had R^2^_GLMN (C, M)_ values of 0.47 and 0.41, slightly higher than the top-performing model (Table 2). Human abundance varied, with the period between 7:00 and 11:00 showing the highest average peak in human maxN. Although the highest maxN per frame was 11 humans at Hithadhoo Corner in April 2023, 98.5% of photos contained 0. Between sites, the mean number of humans per photo, per hour for Hithadhoo Corner, Fushi Kandu, and Fonadhoo Beyru was 0.04 ± 0.40, 0.03 ± 0.42, and 0.03 ± 0.32, respectively. Boduhuraa Beyru had the fewest photos per hour with humans present, with a mean of 0.01 ± 0.12. However, the inclusion of human abundance in the model increased model complexity, which was penalised by the BIC test, indicating that it did not offer improved explanatory power over the top-performing model.

The inclusion of current speed and heading in a subset data frame for GLMM 2.2 yielded contradictory results for AIC_c_ and BIC (Table 2). Three models were within two AIC_c_ units of the top-performing model, two of which included the current heading and none of the current speed. However, BIC showed full support for the null model, with those models supported by AIC_c_ having considerably increased BIC scores. These models should be inferred with caution, as such large discrepancies between AIC_c_ and BIC shown in Table 2 can indicate a lack of explanatory power from the addition of the current heading.

### Re-sighting Patterns

Repeated reidentification of *M. alfredi* provides powerful insights into residency patterns and social demographics. Successful identification was possible on 629 occasions, involving 81 unique individuals, representing 51.59% of *M. alfredi* identified in Laamu Atoll (n = 157, [60]). Ten of these individuals were sighted only once, while 45 were sighted ≥ 5 times. One individual (MV-MA-3004) was captured 33 times. Overall, 24.2% of estimated *M. alfredi* were successfully identified. However, image quality varied across sites, with the few deployments at Fonadhoo Beyru being particularly successful, in which 60.5% (n = 107) of estimated *M. alfredi* were identified, whereas at Fushi Kandu the opposite was true, with only 12.9% identified (n = 4).

Sighting interval times varied spatiotemporally (Fig 6). The mean sighting interval for *M. alfredi* sighted on multiple occasions was one year (352.65 ± 260.48 days). The mean number of days between sightings for individuals sighted multiple times was 62.32 ± 4.04 days (n = 71), with MV-MA-2552 having the longest interval between sightings (519 days). Mean intervals between sightings tended to decrease as the number of sightings increased. The mean sighting interval for a second sighting was 100.36 ± 121.74 days. An individual’s sighting interval decreased to 28.62 ± 30.5 days by their 9^th^ sighting (n = 29), and the lowest interval, which applied to 10 individuals, was 12.3 ± 13.49 days on their 20^th^ sighting, suggesting some individuals stayed in proximity to the cleaning stations. The longest interval from the first sighting on 5^th^ January 2021 to the final sighting on 18^th^ May 2023 was a period of 747 days (MV-MA-3754) and consisted of 23 sightings at Hithadhoo Corner only. In contrast, MV-MA-3755 was sighted only twice between 9^th^ June 2021 and 10^th^ June 2021 at Hithadhoo Corner and has not been sighted since. Intra-specific variation in periodicity was clear. For example, MV-MA-2900 and MV-MA-2927 appear to occupy cleaning sites more seasonally, whereas MV-MA-2551 and MV-MA-2862 exhibit more consistent annual residency patterns (Fig 6). Multiple-site use was apparent in 53 individuals (74.64%), with evident intra-atoll migrations occurring over short time frames (Fig 6). For example, MV-MA-2862 was sighted at Hithadhoo Corner on the 8^th^ of October 2022 and two days later at Fushi Kandu, a ∼30km intra-atoll migration (Fig 6).

**Fig 6.**
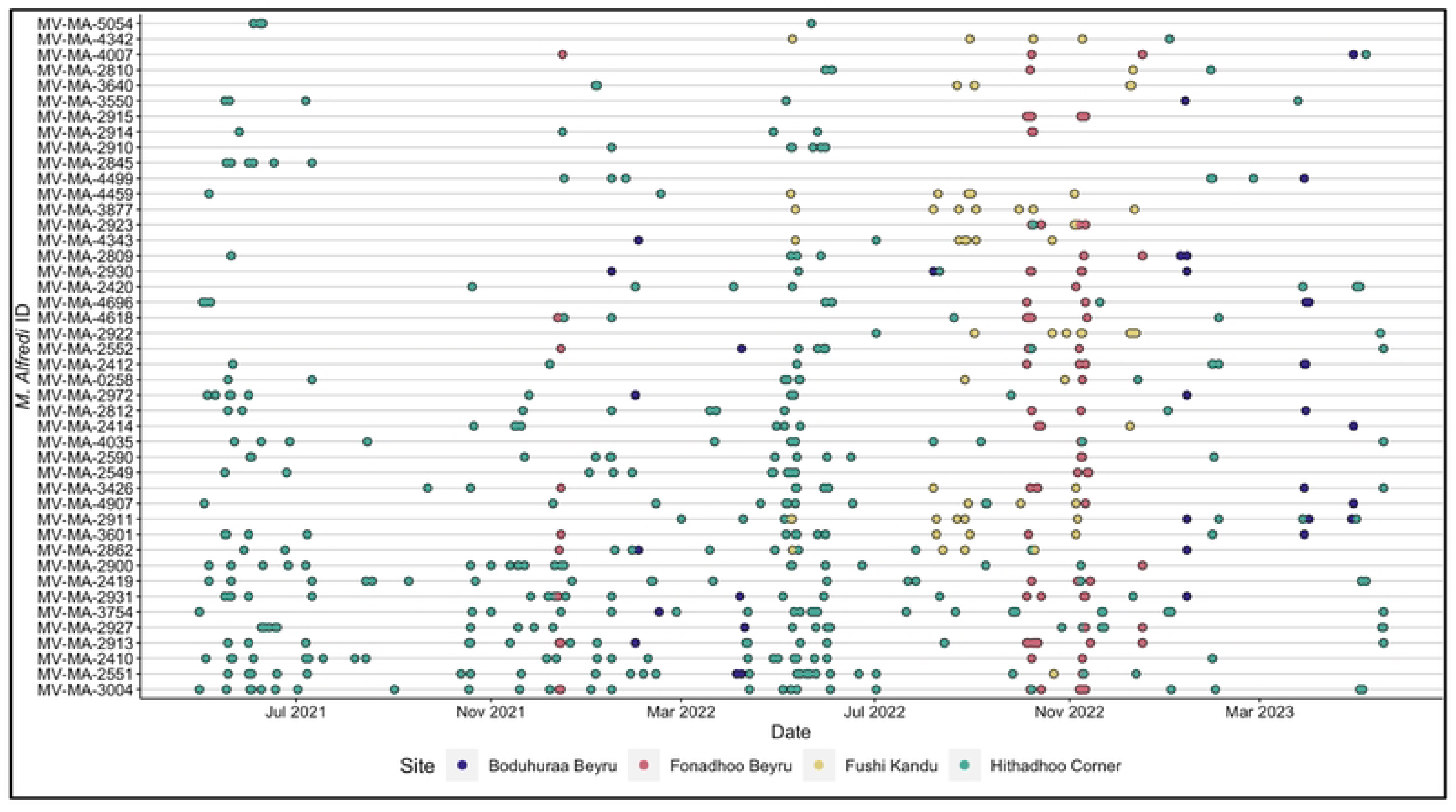
Sightings of previously identified Mobula alfredi at all four cleaning sites. Sightings of individuals from the Manta Trust’s identification catalogue, recorded by remote underwater photo systems throughout the survey period, for those sighted five or more times.

## Discussion

### Drivers of Abundance

#### Annual Productivity

Using novel, low-cost methods to survey a socioeconomically important *M. alfredi* population, this study advances our understanding of cleaning-site use and the drivers of abundance around Laamu Atoll, Maldives. The world’s largest monsoonal variation [73] affected *M. alfredi* abundance at Laamu Atoll’s cleaning stations, depending on their location around the atoll. Previous studies from elsewhere in the Maldives Archipelago are in support of monsoonal seasons affecting *M. alfredi* abundance [7,38,54]. Monsoonal trade winds drive leeward upwellings, increasing productivity [54,73,74], measured here as the chlorophyll-a concentration, which also positively correlated with *M. alfredi* cleaning station abundances. Despite the ultimate and proximate drivers of cleaning station use by *M. alfredi* remaining unclear, an increase in cleaning site attendance during periods of elevated local productivity has been explained by a reduction in the foraging time required for individuals within their home range, leaving more time for cleaning and social activities, which are known to occur frequently at these sites [43,75]. Additionally, an increased need for gill plate cleaning and post-foraging thermoregulation from periods spent in deep, cool waters has been inferred for other mobulid species [76–78]. The oceanographic characteristics of Laamu Atoll, which is surrounded by deep water and productive upwelling, likely play a role in the behaviours identified in this study. Fushi Kandu is in the lee of the atoll during the southeast monsoon, with increased productivity and inferred abundant nearby foraging opportunities potentially contributing to the seasonal increase in presence here.

Throughout Laamu Atoll, *M. alfredi’s* presence at the study sites declined during the Northeast Monsoon. The productivity of the Northeast Monsoon is considered lower than that of the Southwest Monsoon due to weaker prevailing winds, which lead to less intense upwelling and, in turn, comparatively lower productivity [79,80]. *M. alfredi* could travel further to satisfy foraging requirements, and demonstrate dietary plasticity, opting to feed on demersal zooplankton over pelagic zooplankton and frequent cleaning sites less from a reduction in cold water foraging and reduced reproductive opportunities. The results of this study indicate a reduced effect of seasonal movements within Laamu Atoll compared with the cross-national migration observed further north [10,54]. The influence of the South Asian Monsoon diminishes southward of the Indian subcontinent, thereby increasing the relative importance of equatorial current systems, such as the South Equatorial Current, in shaping atoll-scale circulation and ecosystem dynamics [81,82]. This reduction in the South Asian Monsoon has been suggested to be an influential factor in the *M. alfredi* population ecology in the Maldives’ southernmost atoll, Addu, where they are found year-round, similarly to Laamu Atoll, exhibiting strong fidelity to channel-cleaning sites [83]. This shift in the impact of the South Asian Monsoon may be a causal factor in the smaller size and reduced migratory capacity of M. alfredi subpopulations in southern atolls. An annual high abundance here may indicate the importance of this site as a refuge from the weakening South Asian Monsoon [49], as well as the quality of its cleaning opportunities. The Southwest Monsoon is of particular importance in the Maldives, lasting for six months annually and consequently boasting a prolonged period of productivity [7]. Stevens [7] also indicates the importance of the monsoon retreat for social interactions and reproductive behaviour at cleaning sites, where observed courtship displays increased in March, October, and November, whereby high fitness from a productive season lends to an increase in reproductive effort [54]. The high chlorophyll-a and increased abundance of *M. alfredi* around cleaning sites in the months before November in Laamu Atoll may suggest these theories apply here. Continued deployment of RUPs will strengthen conclusions drawn from Boduhuraa Beyru and Fonadhoo Beyru, although the high presence at Fonadhoo Beyru may further support the importance of the Gaadhoo-Hithadhoo Channel for this *M. alfredi* population.

#### Diurnal Visitation

The analysis of fine-scale parameters raised suggestions on diurnal visitation. *Mobula alfredi’s* presence in our study supports others that provide holistic, high-resolution temporal understanding (acoustic: [14,84]; satellite telemetry: [16]), as similarly demonstrated in a related species, *Mobula birostris* (acoustic: [34], satellite telemetry: [9]). Peak abundance in Laamu Atoll was between 09:00 and 10:00, which is comparable with the Chagos Archipelago [8], Indonesia [85], Mozambique [86], and Seychelles [87], where presence peaked between mid-morning and noon, suggested to be driven by cleaning quality and foraging opportunities. Although this study lacks nocturnal identification of *M. alfredi* presence and absence, the similarity of diel cleaning site presence to that of others may suggest that spatiotemporal abundance is driven by similar factors. Furthermore, the crepuscular movement of benthic zooplankton and subsequent foraging could explain the lack of cleaning station attendance at dawn and dusk. Cleaning services are optimised during the day when *Labroides spp.* are most active, and visual interspecific cues from clients to initiate cleaning can be reciprocated [88–90]. Additionally, *M. alfredi* express dietary plasticity, surface feeding during the day, observed infrequently in Laamu Atoll [60], and deep foraging at night on emerging zooplankton, identified elsewhere through stable isotope analysis [91] and multi-function data loggers [55]. The opportunistic use of cleaning sites during this aforementioned change in foraging has been suggested [91] and could help explain the patterns observed here.

The inclusion of the moon state in GLM 1.1 supports theories of forage-driven cleaning site presence [92]. Increases in abundance during low-illumination moon states are well documented [44]; elsewhere: [10,43,85,93]. During darker nights, zooplankton diel vertical migration is more intense, and throughout the full moon, zooplankton predation-grazing trade-off thresholds remain deeper [94]. Reduced *M. alfredi* abundances at cleaning sites around Laamu Atoll during the full moon may necessitate longer and/or deeper foraging, with the energetic cost of returning to clean sites too high [8,34]. Alternatively, *M. alfredi* may seek out other foraging locations, such as atoll lagoons, thereby negating the need to retreat to outer-reef cleaning sites for thermoregulatory and predatory avoidance [95].

The absence of an effect of tide and current in our study may indicate adequate foraging opportunities near the atoll. It is possible that reliance on fine-scale environmental factors to increase patch densities, as populations in Chagos do [64], is absent in Laamu Atoll. Other studies support the need to accurately quantify current, as it affects cleaner wrasse activity, sediment suspension and cleaning energy expenditure [43,64,90]. The short deployments of current meters at Hithadhoo Corner during the Northeast Monsoon, when presence was lowest, may have contributed to inconclusive data, so continued effort is needed here. Furthermore, the limited effect of temperature on presence was expected, given the narrow range of temperatures recorded and previous studies indicating that temperature is a less influential driver of *M. alfredi* presence at cleaning sites [45]. Considering that temperature has recently been described as significant in determining *M. alfredi*’s presence at foraging sites elsewhere in the Maldives [49], it is necessary to identify individual temperature profiles or deploy offshore and outer-reef sensors to further our findings [36].

### Re-sighting Patterns

Sighting intervals differed among individuals, as expected due to differences in demographic, sex, energetic, and thermoregulatory demands [8,85,87,96]. Year-round sightings were observed for some *M. alfredi* at multiple sites, while others appeared to exhibit cleaning site fidelity, behaviours also observed in Baa Atoll, Maldives [44] and Raja Ampat, Indonesia [33]. This is further supported by a mean re-sighting interval of 62.3 days, which was negatively correlated with the number of times an individual was sighted. Compared to northern atolls, which demonstrate high degrees of connectedness [20], only 17.2% (n = 157) of individuals ever recorded in Laamu Atoll have been identified elsewhere [60]. Resighting of individuals four times or fewer occurred 44% of the time (n = 31), which is conservative given that RUP systems fall short of determining absolute absence and had an identification success rate of 24%. Compared with observational surveys, identification success rates typically exceed 90% and enable the identification of annual multi-demographic residencies, copulation, and pregnancies [7].

A variety of factors may be driving these differences in re-sightings, including bathymetry and atoll isolation [10,11]. Life history needs of the small Laamu Atoll *M. alfredi* population may be satisfied by the island mass effect theory, thus removing their requirement to migrate to other atolls [34,54]. Local bathymetry around Laamu exceeds ∼1500 m to the north of the atoll and ∼2500 m to the south, within ∼5 nautical miles. The predation risk associated with deeper water may be too high [12], especially as in-shore conditions around Laamu could be satisfactory year-round. A genetic study on *M. alfredi* in Hawaii could provide comparisons to these theories. Whitney *et al.* [12] concluded that two islands 46 km apart, where individual movement has never been recorded, have genetically distinct populations, due to a 3000 m channel and satisfactory foraging opportunities to support isolated populations. However, despite the geographic similarities between Laamu Atoll and Hawaii, Hosegood [97] concluded that, based on five individuals from Laamu Atoll and 18 from Baa and Raa Atolls, inter-atoll connectedness was sufficient to ensure genetic homogeneity in the region. Laamu Atoll’s geographical isolation, compared with Hawaii’s [12], and individual variation in residency (Fig 6) may indicate partial residency. While past studies have shown mesoscale fidelity [93] and long-distance migrations [55,98,99], our research raises questions about Laamu forming discrete sub-populations, similar to Hawaii [12] and Raja Ampat, Indonesia [33], within globally recognised populations [13].

### Study Limitations

The RUP deployments showed a trade-off between increased survey effort and reduced data quality compared with SCUBA surveys, which dominate these sites in Laamu Atoll. Stevens [7] encountered a comparable 2.7 *M. alfredi* per survey day whilst using SCUBA, compared to the average 3.5 presented here. Remote underwater photo systems are low-cost, non-invasive, easy to deploy and capture unbiased snapshots of time. However, they rely on long-term observational databases or more informative footage from remote underwater video systems to supplement their findings on individual identification. In contrast to the RUPs used here, SCUBA surveys in Laamu Atoll are also able to comment on *M. alfredi* behaviour, achieve a ∼74% individual identification rate, and identify sex, maturity, reproductive state, and injuries [60]. No published literature has used RUPs exclusively to identify the presence of M. alfredi at cleaning sites. Buschmann *et al.* [86] and this study show how valuable remote observations can be as an additional technique to use alongside observational studies. Combining methodologies, such as opportunistic in-water surveys and RUPs, improves our understanding of *M. alfredi’*s spatiotemporal ecology, importance, and functioning, and provides clear avenues for future research and conservation priorities [15,75,100].

This study has some drawbacks, as do many photographic-based studies [15]. Sighting events defined by an arbitrary interval of 10 minutes inflate pseudo-replication and, therefore, the estimated abundance. Accurate identification of the intervals between cleaning events using acoustic tags would be powerful. In opposition to the estimated abundance, maxN is considered a definite and conservative measure of abundance [101], but neither count provides insight into absence, a shortfall of many observational techniques, something that 360° cameras may solve. Time of first arrival or maximum number of individuals (maxIND) and indices of residency [9] were all infeasible, given that the systems are passive. Here, 24.2% of individuals were successfully identified, which is low compared to the near 100% success rates reported for SCUBA/snorkel photographs, remote underwater video, and acoustic telemetry [7,8,42]. This limits direct statistical comparison of the Laamu Atoll cleaning sites with others, reducing the credibility of some inferences from the results. Additionally, the narrow GoPro field of view creates a trade-off; either capturing the whole cleaning site or high-resolution branchial spot images. Systems were placed in favour of quality branchial spot photographs, reducing the quantity of the cleaning site observed. Issues with parallax errors, low-quality images, biofouling, floating debris and partial branchial photographs further jeopardised identification.

### Mobula alfredi Conservation in Laamu Atoll

*Mobula alfredi* is one of the least fecund species in the world and is listed as Vulnerable to extinction on the IUCN’s Red List [5,13]. Fishing is their greatest threat [102,103]. Regional law prohibits the direct capture of manta rays within the Maldives [51]. Most of the time, *M. alfredi* are absent from cleaning stations, highlighted by the median sighting duration of four minutes. Observational studies also support the varied site use of *M. alfredi* and indicate that protection must extend beyond cleaning sites alone [36]. A national *M. alfredi* management plan is needed to effectively protect the Maldives population and their critical habitats from the plethora of threats they face.

Rises in sea surface temperatures, resulting in increased Maldivian coral bleaching alerts [104] and a 20% reduction in zooplankton over the last half-century [105], as well as land reclamation, which is common throughout the Maldives [106], highlight the need for urgent local and global conservation efforts. Positively, anthropogenic presence did not affect abundance; a result also found by Buschmann *et al.* [86]. RUPs infrequently captured breaches of the Manta Trust’s code of conduct despite threats from tourism generally remaining low at these study sites. The economic necessity and growth of tourism, but the observed negative effects on manta rays in the Maldives elsewhere [20,26,50], provide a warning that effective marine protection needs to be gazetted. Laamu Atoll’s *M. alfredi* population is smaller than others identified in the Maldives [7] and shows signs of partial isolation. Therefore, vulnerability and inter-individual reliance are amplified, and there is a clear need for species- and atoll-specific legislation to protect them.

Laamu Atoll’s governance currently provides no additional protection for the resident *M. alfredi* population, despite its strategic location. Hithadhoo Island is the first Maldivian community to propose managing its own Community Conservation Area, which includes Hithadhoo Corner, a newly designated Important Shark and Ray Area [29] (Fig 1). Maldives Resilient Reefs, a local non-governmental organisation affiliated with BLUE Marine Foundation, is currently drafting the management plan in consultation with the residents. The importance of community engagement is clear for long-term, sustainable protection [28]. Our current understanding of the spatiotemporal site use of Laamu Atoll’s *M. alfredi* is insufficient to propose an informed network of connected MPAs. Therefore, a highly protected, atoll-wide MPA should be considered, especially given our results demonstrate the importance of the atoll for consistent annual *M. alfredi* presence. Additionally, implementing larger MPAs is operationally and economically effective [107]. As with the Chagos archipelago [108], such an MPA would directly protect the *M. alfredi* and their associated habitats [109]. Efforts to identify other key aggregation areas will further MPA planning specificity and ultimately, effectiveness.

Two clear research priorities emerged from our study. *First*: the requirement to identify key *M. alfredi* foraging aggregations. Specific protection must extend beyond cleaning sites to ensure this species’ longevity. Argos-linked satellite tags, such as Wildlife Computers Fastloc [16], could provide accurate insights into the breadth of habitat use, inter-atoll connectivity, and foraging sites. *Second*: To what degree the Laamu Atoll *M. alfredi* population interacts with and is affected by the local fishery. Execution of these research priorities, with continued SCUBA/snorkel-based observations, RUP deployment, and exploration of Laamu Atoll, will improve our understanding.

## Acknowledgements

We thank the Government of the Maldives for permitting us to conduct the research. We would like to thank Six Senses Laamu and Marteyne van Well for hosting the fieldwork component of this study, for their continued financial support, and for facilitating the Eyes on the Reef project. We thank all Maldives Manta Conservation Programme staff, students and volunteers, as well as the wider Manta Trust team (past and present, especially Tam Sawers). Thank you to the team of staff and interns from the Maldives Underwater Initiative (MUI) for their assistance with fieldwork, and to Deep Blue Divers for organising the boat logistics. We would also like to thank the Hithadhoo Council, the Fonadhoo Council, and the Maabaidhoo Council for supporting the research conducted within their jurisdictions. This work has evolved since its submission by the author, BJGP, as their Master of Science thesis to the University of Exeter. For the purpose of open access, the authors have applied a Creative Commons Attribution (CC BY) licence to any Author Accepted Manuscript version arising from this submission

## Notes

### Competing Interest Statement

The authors have declared no competing interest.

